# Lipid-induced polymorphic amyloid fibrils formation by α-synuclein

**DOI:** 10.1101/2021.07.20.453062

**Authors:** Bhanu Pratap Singh, Ryan J Morris, Mathew H Horrocks, Tilo Kunath, Cait E MacPhee

## Abstract

Many proteins that self-assemble into amyloid and amyloid-like fibres can adopt polymorphic forms. These forms have been observed both *in vitro* and *in vivo* and can arise through variations in the steric-zipper interactions between β-sheets, variations in the arrangements between protofilaments, and differences in the number of protofilaments that make up a given fibre class. Different polymorphs which arise from the same precursor molecule not only exhibit different levels of toxicity, but importantly can contribute to different disease conditions. In this work, we show that in the presence of 1,2-dimyristoyl-sn-glycero-3-phospho-L-serine, a highly abundant lipid in the plasma membrane of neurons, the aggregation of α-synuclein is markedly accelerated and yields a diversity of polymorphic forms under identical experimental conditions. This morphological diversity includes thin and curly amyloid fibrils, helical and twisted ribbons, nanotubes and flat sheets. TEM analysis of fibrils sampled from the early stage of the growth phase shows the presence of helical and twisted ribbons, indicating that these morphological variants form at the early stages of aggregation. Total internal reflection fluorescence microscopy (TIRFM) indicated the presence of lipids collocated with the mature fibrils. This finding has important implication as the presence of α-synuclein with co-localized high lipid content has been reported in Lewy bodies, the pathological hallmark of Parkinson’s disease and other synucleinopathies. Thus, the present study demonstrates that an interface, such as that provided by a lipid membrane, can not only modulate the kinetics of α-synuclein amyloid aggregation but also plays an important role in the formation of morphological variants by incorporating lipid molecules in the process of amyloid fibril formation.

## INTRODUCTION

Protein self-assembly is involved in many diverse biological processes, from maintaining cellular homeostasis to being the causative mechanism of many diseases. Some well-known examples of nature utilising self-assembled protein structures are the cytoskeletal architectures built from actin and tubulin^1,2^, the aggregation of blood fibrinogen into fibrin^3^, the formation of collagen fibres^4^, and the synthesis of spider silk^5^ to name but a few. Under the appropriate conditions, however, many proteins can self-assemble into supramolecular structures known as amyloid fibrils that are associated with more than thirty human diseases including Parkinson’s and Alzheimer’s disease^6^. Apart from their importance in the role they play in disease, recently amyloid fibril structures have been successfully utilized in the field of nanotechnology, bioelectronics, biomedicine and material science due to their exceptional mechanical properties^7,8^. Thus, understanding the factors that control and modulate the processes of protein self-assembly is of fundamental importance and will impact multiple areas of current scientific research.

Amyloid fibre formation has been reported for a large number of proteins, including many proteins that are not known to form amyloid fibrils *in vivo^9^.* Although these amyloid fibre forming proteins are distinctly different in primary amino acid sequence^10^, the shared structural features are strikingly similar. Studies carried out by solid-state NMR, cryo-EM and diffraction methods^10,11^ have enriched our knowledge of the common structure elements of amyloid fibrils. These studies show that amyloid fibrils consist of orderly arrangements of β-sheets and β-strands that are organized parallel and perpendicular to the fibril axis, respectively. Each β-sheet interacts with a neighbouring sheet via “steric-zipper” interactions between amino acid sidechains^12^. Interestingly, it was observed in multiple amyloid fibre forming systems that the β-sheets can pack via different symmetry classes of steric-zipper interactions^13,14^, which results in different conformational variants known as polymorphs. Another factor which can contribute to amyloid fibril polymorphism is the differential packing of protofilaments into mature fibrils, which arises due to differences in lateral contacts between protofilaments^15^. For example, a cryo-EM study of two different polymorphs of tau fibrils (paired helical and straight fibrils) extracted from an Alzheimer’s diseased brain show that these variants have two identical protofilaments but have differential packing arrangement of protofilaments^16^. Such polymorphs have been observed under both *in vitro* and *in vivo* conditions for various proteins. With a growing number of studies, this phenomenon is emerging as a general feature for all amyloid fibre forming proteins proteins^13,17,18^.

Apart from the different steric-zipper interactions that can occur between the β-sheets of a protofibril and the differential packing arrangements of protofilaments into mature fibrils, another factor that contributes to the observed polymorphism is the number of protofilaments that make up the mature fibril^19^. The polymorphic forms of these fibrils possess interesting geometrical features such as helical and twisted architectures, variable widths (10 nm-173nm), and higher persistence lengths which are directly correlated with the protofibril number ^20,21^.

Importantly, different polymorphs arising from the same precursor protein exhibit different level of toxicity and different disease conditions^18,22^. For example, amyloid fibrils isolated from brain cells of Alzheimer’s patients have been found to be morphologically homogenous in a single patient but different from one patient to another^23^ and the different polymorphs of these fibrils formed by the β-amyloid protein display different degrees of toxicity in neuronal cells^24^. Moreover, in addition to differences in cell toxicity in a single cell line, different amyloid fibril polymorphs can also target different tissues and cell lines resulting in distinctive pathologies^25^. In the case of synucleinopathies (e.g. Parkinson’s disease, multiple system atrophy, dementia with Lewy bodies), these conditions are all associated with the formation of α-synuclein (α-syn) amyloid fibrils^26^. However, the amyloid fibril structures presented in these pathologies are different^22^, i.e. these diseases are correlated with different polymorphic forms of α-syn amyloid fibrils. Thus, it is very important to understand and discover the factors that play an important role in the formation of fibril polymorphs.

α-Syn is a small 140 amino acid protein present in the presynaptic terminal of neuronal cells. In its soluble form it presents as an intrinsically disordered protein, whereas in its membrane bound form the N-terminal region acquires an α-helical rich structure^27^. Its membrane bound form has been linked to its putative physiological role in synaptic plasticity, neurotransmission release, assembly and disassembly of the SNARE complex^28^. Interestingly, a recent study showed the importance of native α-syn in the central nervous system for providing innate immunity against RNA virus infection^29^. Importantly, many *in vitro* studies have shown that its interaction with different lipid membranes plays an important role in modulating the kinetics of amyloid fibril formation^27^.

It is well known the lipid composition of the brain changes with the age^30^ and PD is largely age associated pathological condition^31^. Phosphatidylserine in one of the most abundant lipid in the synaptic vesicles^32^ and importantly, it has been shown previously that phosphatidyl serine synthase has an elevated activity in the substantia nigra of patient with Parkinson’s disease^33^. For this reason, we have carried out our investigation with 1,2-dimyristoyl-sn-glycero-3-phospho-L-serine (DMPS) and we have shown that in the presence of DMPS, fulllength human α-syn monomers aggregate into several polymorphic forms. Their morphologies include thin and curly amyloid fibrils, helical and twisted ribbons, nanotubes, and flat sheets. Image analysis of fibril characteristics shows the large variations of polymorph within each class; such a high number and different class of amyloid polymorphs under a single set of environmental condition has hitherto not been reported for any protein system. Time-dependent studies of fibril formation revealed the presence of helical and twisted ribbon variants at the beginning of the growth phase of amyloid fibrils. This observation contradicts previous models that proposed that these high-order polymorphs occur by lateral-association of mature protofilaments at much later stages of fibril assembly^19,20^. Finally, using TIRF microscopy we show evidence for the incorporation of lipid molecules into the mature fibrils. Thus, this study shows that the presence of DMPS not only modulates the kinetics of α-syn amyloid formation, but also can be integrated into the mature fibrils.

## RESULT AND DISCUSSION

### Effect of DMPS vesicles on the kinetics of α-synuclein fibrillization

Full-length human α-syn was expressed recombinantly in *E. coli* and purified as described in methods. Aggregation assays were subsequently performed in the presence or absence of DMPS vesicles under static conditions. The kinetics of amyloid fibril formation was monitored by ThT fluorescence and all experiments were conducted in aqueous solution at pH 7.4 at 30°C. We observed that in the presence of DMPS vesicles the lag time of fibril formation was dependent upon the lipid/protein molar ratio. It was found that increasing the value of this ratio corresponded to a decrease in the lag time (**Fig. 1A**). However, significant variation in the lag time was observed at all ratios tested (0.5, 1, 2, 4 and 8) (**Figs.1 and S2**). The mechanistic details for lag-time variability is not well understood, however, the stochastic nature of nucleation at the molecular level may influence the observed variability at the macroscopic scale^34^. Indeed, α-syn is known to exhibit highly variable self-assembly kinetics between replicates of the same biological sample^35^. These data imply that DMPS vesicles significantly promotes the fibrillization of α-syn, but it does not affect the lag-time variability of protein fibrillization.

**Figure 1.**
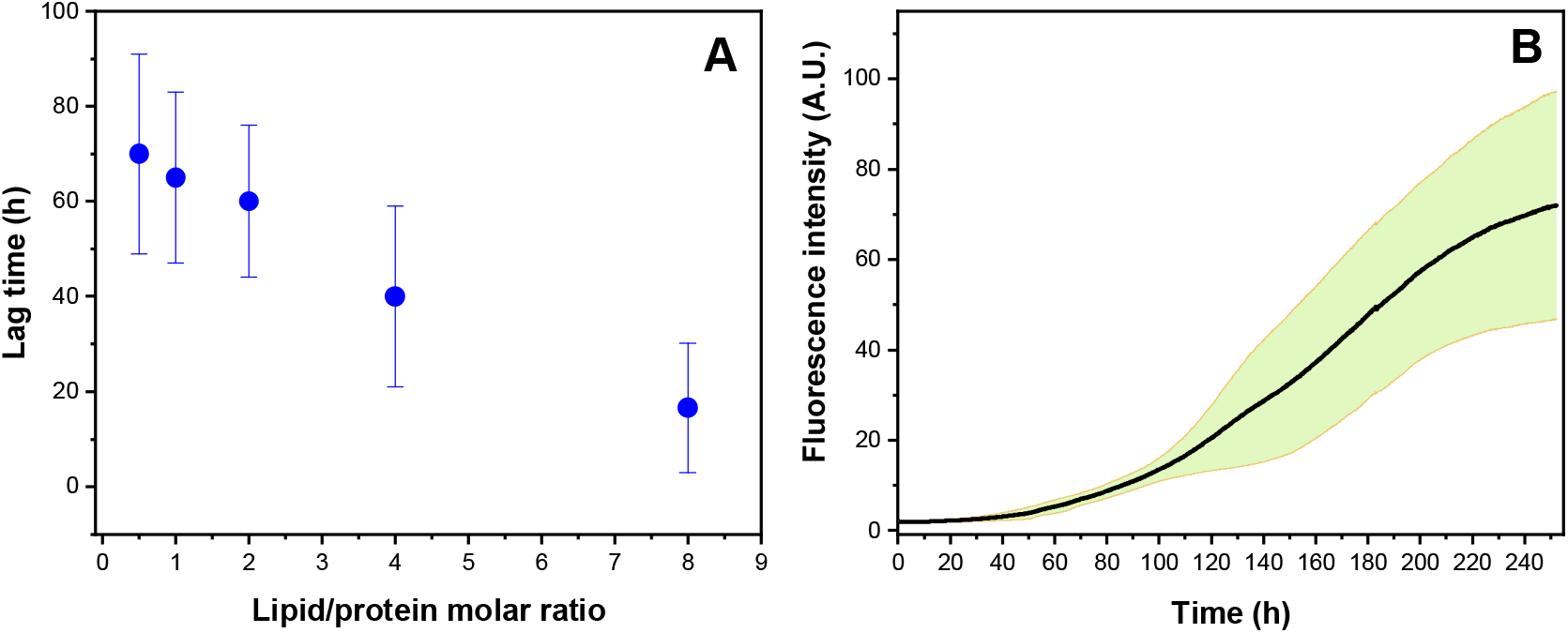
**(A)** Plot of lipid:protein ratio against lag time in amyloid formation by α-syn in the presence of DMPS vesicles, n =8, error bars are standard deviation **(B)** Kinetics of amyloid formation by α-syn (50 μM) and DMPS at a lipid:protein molar ratio of 8, as monitored by ThT fluorescence. Shaded green region is the standard deviation.

α-Syn is known to interact with various lipids which can differentially modulate the kinetics of fibril formation, either by promoting or inhibiting the process. The lipid/protein ratio is an important factor for the observed differential effects on the kinetics of α-syn aggregation. When this ratio is relatively low it promotes the process. At higher ratios, however, the process is inhibited as there is a depletion in free monomer in solution as α-syn preferentially binds to lipid membrane^32^. For our initial experiments investigating the kinetics of assembly in the presence or absence of DMPS, we chose a physiological concentration of α-syn protein (50 μM)^27,36^ and a lipid/protein molar ratio (8) that minimised the lag-time for fibril-formation^32^. We found that at this ratio the lag time of reaction varies from about 4h to 45h (**Fig. 1B**) with a mean value of about 16 h (**Fig. 1A**). In the absence of DMPS vesicles only a very small increase in ThT signal was observed at the same protein concentration (**Fig. S3**), however, increasing the protein concentration to 100 μM showed a significant increase in signal. The reaction was allowed to progress for 10 days and the samples that reached the plateau phase were selected for morphological characterisation by TEM.

### Morphological study of amyloid fibrils by TEM

TEM imaging of the sample in the absence of DMPS vesicles shows the presence of small oligomeric aggregates when the concentration of the incubated sample was 50 μM (**Fig. 2A**). However, we observed fibrils with a rod-like morphology when the concentration of protein was 100 μM (**Figs. 2B and Figs. 2C**). In contrast, TEM images of the amyloid fibrils of α-syn formed in the presence of the DMPS vesicles exhibited a large diversity of morphologies (Figure 2D - Figure 2I). These images show the existence of different morphological structures of amyloid fibrils, including thin and curly amyloid fibrils without any clear periodicity (green arrows, Fig. 2D), helical (blue arrows, Fig. 2F) and twisted ribbons (red arrows, Figs. 2E & Figs. 2F), nanotubes (yellow arrows, Figs. 2G & Figs. 2H), and flat sheets (purple arrow, Fig. 2I). While thin and curly amyloid fibrils appear to be the earliest stable fibril form, the other forms, (twisted and helical ribbons, nanotubes and flat sheets) are formed by the high-order association of thin and curly amyloid fibrils. The characteristic feature of twisted ribbons is the presence of anisotropic cross section and saddle-like (Gaussian) curvature, whereas helical ribbons can be wrapped around a hypothetical cylinder of a finite radius and characterized by defined mean curvature associated with bending but zero Gaussian curvature^19,37^. Although twisted and helical ribbons both consist of multiple protofilaments, the transition from twisted ribbons to helical ribbons takes place when the number of protofilaments reaches a critical number (parameterized by the width to thickness ratio). When this ratio is low, twisted polymorphs are energetically more favoured, whereas at high ratios helical polymorphs are energetically more stable^19,37,38^. Further, helical ribbons can transform into nanotubes, which is a consequence of eliminating line tension associated with the exterior protofilaments of a fibril^19,39^. Flat sheets can form by the assembly of multiple parallel protofilaments and these structures have a higher stiffness than helical ribbons^37,40^.

**Figure 2.**
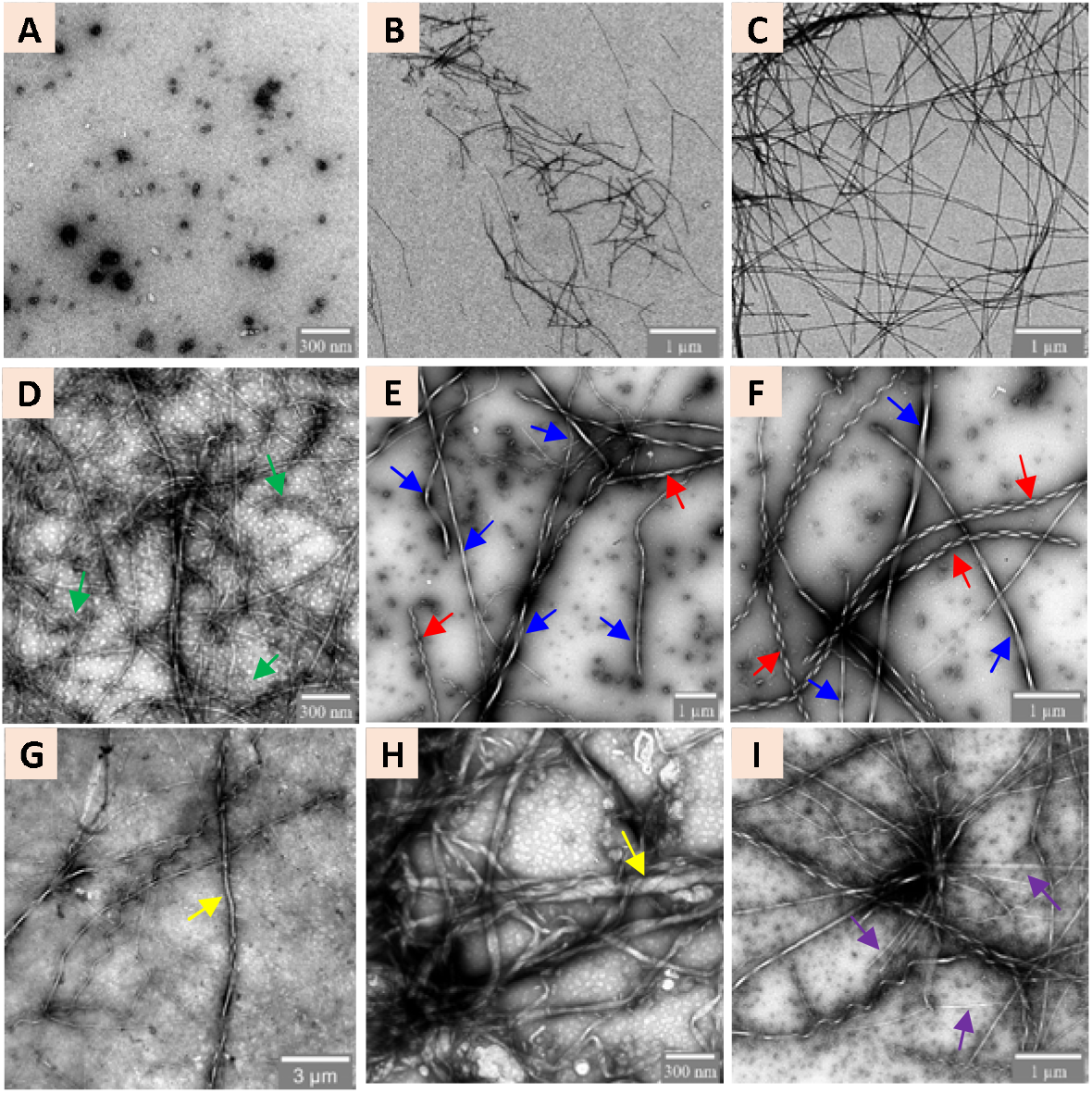
TEM imaging. Polymorphs of α-syn (50 μM, 100 μM) in the absence and presence of DMPS vesicles (400 μM). **(A)** 50 μM of α-syn in absence of DMPS vesicles shows formation of small oligomers. **(B-C)** 100 μM of α-syn in the absence of DMPS vesicles shows presence of rod-like fibrils. **(D)** Thin and curly amyloid fibrils marked with (green arrow) **(E-F)** twisted (blue arrow) and helical (red arrow) ribbon. **(G-H)** nanotube (yellow arrow) and **(I)** Presence of flat sheet (purple arrow). Images show gross morphological features; note the variation in scale bar across images.

These results clearly demonstrate that in the presence of DMPS vesicles, α-syn aggregates into diverse polymorphic forms. Compared to previous data, the presence of such a large number of polymorphic variants formed under identical environmental conditions has not been reported before for any protein. Although experimental data for the kinetics of α-syn aggregation has been reported in the presence of a variety of lipids, the morphological details are limited to only a few studies^27,32,48–52,35,41–47^. It was previously reported that in the presence of phospholipid vesicles composed of 1,2-dipalmitoyl-sn-glycero-3-phosphate (PA) and 1,2-dipalmitoyl-sn-glycero-3-phospho-choline (PC) there was no effect on the morphology of the fibrils as compared to in the absence of these vesicles^52^. Surprisingly, in the presence of small unilamellar vesicles composed of DMPS or DLPS, α-syn has been shown to form small spherical fibrils coated with lipid vesicles and thin fibrils attached to vesicles^32,49^. In another recent work with low lipid: protein ratio, it has been shown α-syn forms many polymorphic variants which includes helical and twisted morphology^53^. However, the kinetic data of amyloid fibril formation is very different than our observation _ their data shows very short elongation phase compared to our data. Apart from difference in the kinetic data in our study the maximum value of helical pitch of fibrils is much higher (~18 times). Further, they have observed the presence of these high order aggregates only after long incubation at plateau phase, whereas we show presence of polymorphic forms from the early growth phase. These results have been discussed in more detail in the later section. It is important to note that in the previous study with DMPS vesicles, in their method for fibril preparation, the reaction was carried out at pH 6.4 in a phosphate buffer^32^. The presence of phosphate in the reaction solution in addition to variations in pH has been reported to have a strong effect on amyloid fibril morphology^54,55^, which can cause proteins to adopt entire different conformation and resulting into different polymorphs. Our observation of a large spectrum of polymorphic forms of amyloid fibrils formed in the presence of DMPS differs from these previous reports. All things considered, it appears that many factors, including lipid:protein ratio, the chemistry of the lipid head group and solution conditions (e.g. pH and co-solutes) can influence the formation of polymorphic forms of α-syn fibrils.

### In the presence of DMPS vesicles α-syn does not aggregate via a helical intermediate

It is well established that in the presence of interfaces such as anionic lipids, cardiolipin, poly unsaturated fatty acids, detergent and trifluoroethanol, the secondary structure of α-syn changes from random coil to an α-helical form^27^. Conditions promoting such structural changes are often associated with an increase in the propensity to form amyloid fibrils^56^. Although the structural and conformational dynamics of membrane bound and nonaggregated states of α-syn have been well characterized^57^, it is not well understood if the membrane-bound protein molecules are on the aggregation pathway. Importantly, a study carried out in the presence of sodium dodecyl sulfate (SDS) micelles _ a well characterized phospholipid mimetic-had shown that α-syn can adopt three well-defined equilibrium states (unfolded, high degree of α-helical structure, and an intermediate state), depending upon the micelle concentration^58^. Similarly, in the presence of an organic solvent such as simple and fluorinated alcohol, a multiphasic structural transition of α-syn has been observed^59–60^. Furthermore, the observed transition phases differed for different solvents. A common feature for all of these studies, however, is that all were characterized by the presence of a partially folded intermediate at a relatively low concentration. Importantly, these intermediate species were found to higher propensity to aggregate. To understand if a helical intermediate is involved in the fibril formation of α-syn in the presence of DMPS vesicles, we performed circular dichroism (CD) spectroscopy at different DMPS/α-syn molar ratios from a value 0 to 30 **(Fig 3A)**. In the absence of DMPS vesicles, CD spectra were flat except for minimum at ~198 nm, which is characteristic of a random coil configuration. As we increased the ratio of DMPS/α-syn, we observed a significant change in the CD spectra. Minima appeared at values of 222 and 208 nm which is indicative of the formation of an α-helical conformation. A plot of ellipticity at 222 nm (θ_222_) and 198 nm (θ_198_) shows a linear trend across all lipid/protein ratios. **(Fig 3B)**. This result is consistent with a two-state transition model and shows that in the presence of DMPS vesicles, α-syn aggregation does not involve a helical intermediate. It is important to note that the results presented here are physiologically more relevant compared to similar studies with SDS micelles or organic solvents since DMPS is one of the most abundant components of synaptic vesicle membranes^32^ and its elevated level has been linked to PD^33^.

**Figure 3.**
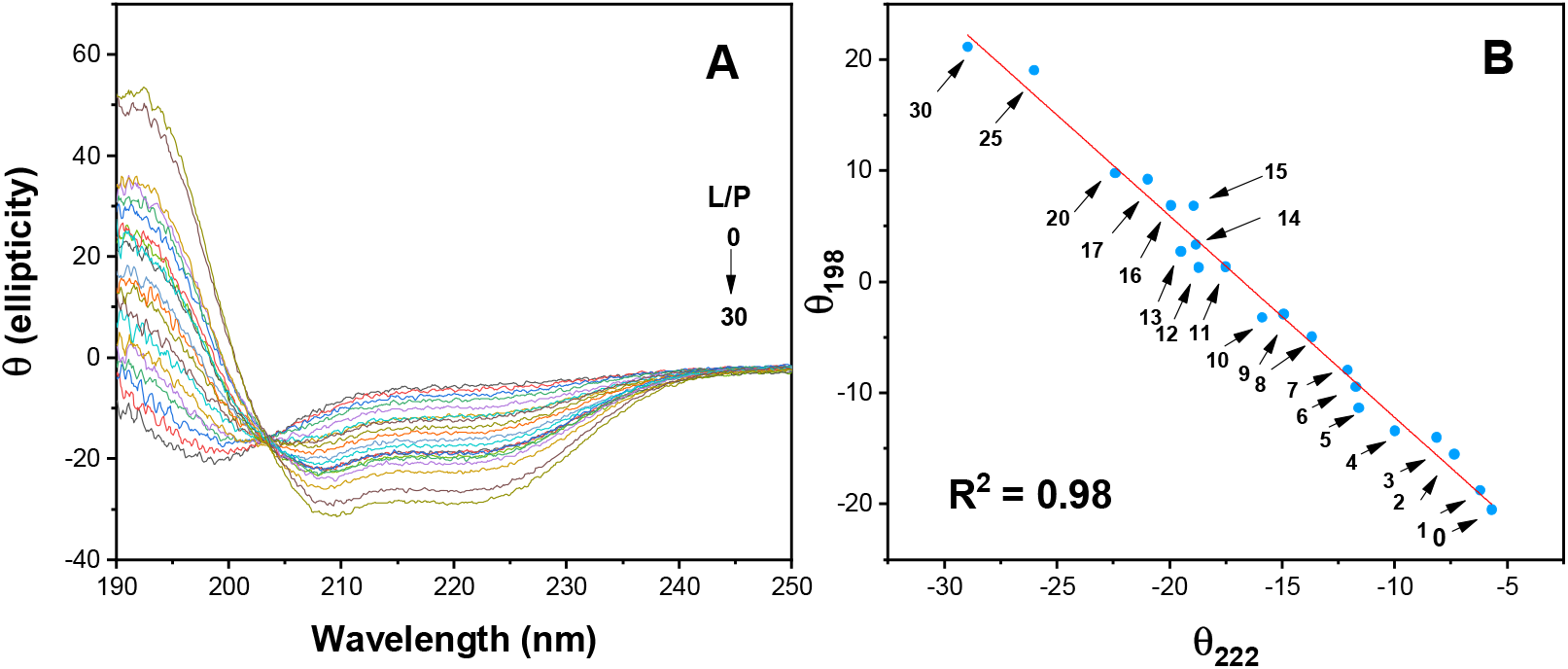
**(A)** Far-UV CD spectra of 10 micromolar α-syn alone and in presence of 10, 20, 30, 40, 50, 60, 70, 80, 90, 100, 110, 120, 130, 140, 150, 160, 170, 200, 250 and 300 micromolar DMPS vesicles. **(B)** Plot of ellipticity at 222 nm (θ_222_) against ellipticity at 198 nm (θ_198_), the numbers denote the lipid/protein ratio across different points.

### Quantitative analysis of amyloid fibrils

As the morphology of α-syn fibrils in the presence of DMPS vesicles was found to be significantly different from those in the absence of DMPS vesicles, we performed quantitative analysis of TEM images to extract the structural characteristics (e.g. length, width) of the polymorphic fibrils. To this end, we measured the width and contour length of 900 and 1200 fibrils, respectively, formed in both the absence and presence of DMPS. A comparison of fibril length and width is shown in **Fig. 4**. The results obtained from this analysis show that in the presence of DMPS vesicles, the average fibril length is 10.8 ± 7.5 μm with a minimum and maximum value of 0.8 and 43.2 μm, respectively. In comparison, fibrils formed in the absence of DMPS were measured to have an average length of 2.6 ± 2.6 μm with a minimum and a maximum length of 0.2 and 15.8 μm, respectively (**Fig. 4A**). Thus, fibrils formed in the presence of DMPS vesicles are much longer compared to those that were formed without. Moreover, such long fibril lengths have not, to the best of our knowledge, been previously reported for α-syn. Analysis of fibril width shows that in the presence of DMPS vesicles, the average width is 30.4 nm with a minimum and maximum of 3 and 157 nm, respectively. On the other hand, fibrils formed in the absence of DMPS have an average width of 16.3 nm, with a minimum and maximum value of 7 and 33 nm, respectively (**Fig. 4B**). For fibrils formed in the absence of DMPS, the width of the fibril remains constant for its entire length. In contrast, the width of fibrils generated in the presence of DMPS shows a constant width for thin and curly fibrils and for flat sheets. However, helical and twisted ribbons possess a periodic modulation in the width as a function of fibril length.

**Figure 4.**
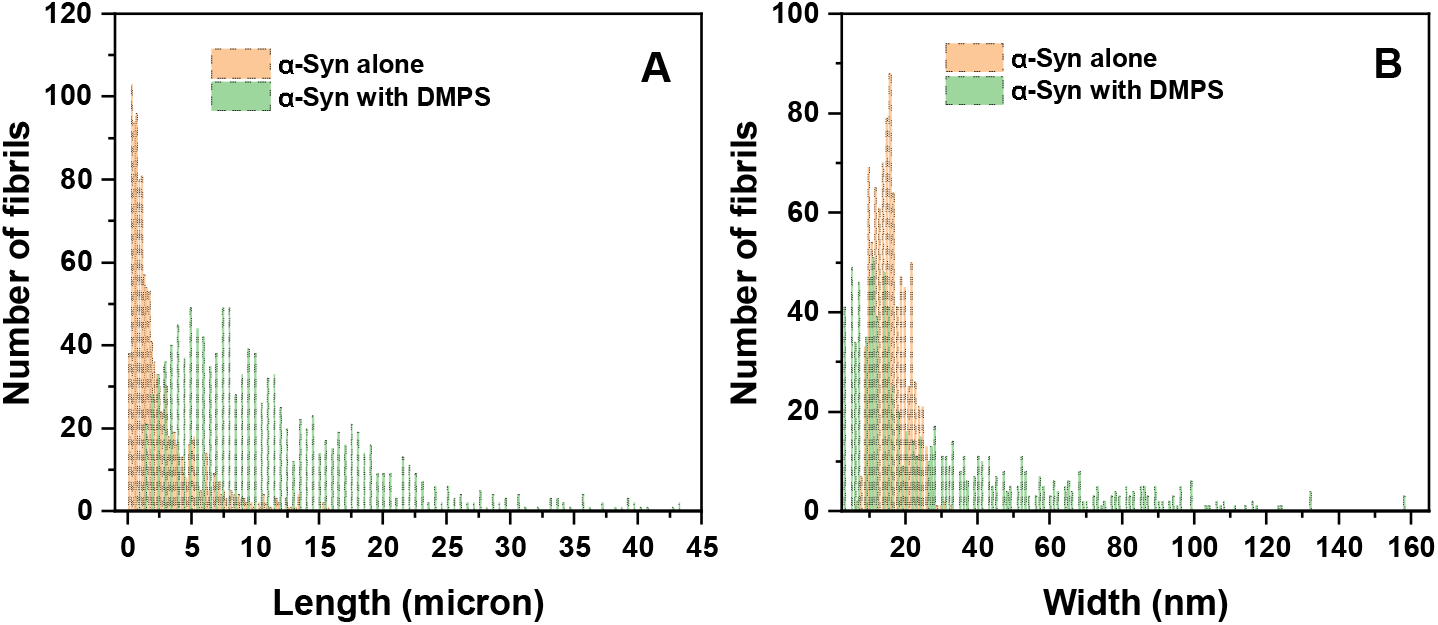
Quantitative analysis: **(A)** Distribution of contour length of α-syn fibrils. **(B)** Distribution of fibril width of α-syn. These results show that in the presence of DMPS the contour length and width of the fibrils is much higher than in its absence.

It has been demonstrated that shorter peptides have a higher tendency than larger proteins to form larger fibril assemblies consisting of laminated β-sheets which arise from the lateral association of protofibrils^61^. Most studies that report the presence of amyloid fibril polymorphs that are comprised of a large numbers of protofibrils have typically been studied in short-peptide systems - either synthetic peptides or fragments of amyloid forming proteins^14,15,40,61–66^. The major drawback of studying small peptides is that in these studies, fibril-formation occurs via homotypic-interactions^14^. In contrast, structural studies of fulllength fibrils at atomic length scales show these interactions are composed of 25-70 residue heterotypic interactions^16,67,68^. For example, the 11-aa peptide of the NAC region of α-syn makes a homotypic steric zipper structure, whereas the same region in the full-length protein bends twice and form an S-shaped heterotypic zipper^68,69^. Although many full-length proteins have been shown to form polymorphic forms, the number of protofibrils involved in these polymorphs is limited to only a few^13,38,54,70–73^. Apart from α-syn, only β-lactoglobulin and lysozyme have been reported to form higher order polymorphic assemblies of full-length protein. For lysozyme it was shown to form both helical and twisted ribbons whereas beta-lactoglobulin can only form helical fibrils^20,72^. For polymorphs consisting of high numbers of protofibrils, it was found that these fibrils were intrinsically unstable and degrade into smaller fragments. The studies in these structures were observed were performed at low pH (2) and high temperature (90 °C); at lower temperatures the same proteins are unable to form higher-order assemblies of protofibrils. The findings described here demonstrate the existence of a large spectrum of polymorphic forms of amyloid fibrils produced at more physiological conditions (both temperature and pH). The fibrils we observe vary widely in their physical characteristics and importantly, are highly stable (**Fig 2, Fig 4 and S4**).

Helical pitch and width are important parameters often used to identify the polymorphs of amyloid fibrils^21^. A helical pitch is defined as the distance along the fibril length that it takes for the helix to complete one 360° turn. **Figure 5** shows the relationship between helical pitch and fibril width for multiple helical fibrils. We found that the helical pitch ranged from 245 to 3373 nm and widths spanned from 15 to 157 nm. Moreover, we found that the helical pitch is linearly correlated with the width of the fibrils measured. Previous work on other polymorphic amyloid fibril systems demonstrated a similar relationship^20,72^. In those works, they found a very strong linear relationship with little variation from the linear model whereas we observe more variation. One interpretation of this finding is that this may imply there are different structural arrangements or interactions occurring between the constituent protofibrils which gives rise to the structural variations we see. However, more work needs to be done in order to confirm this hypothesis.

**Figure 5.**
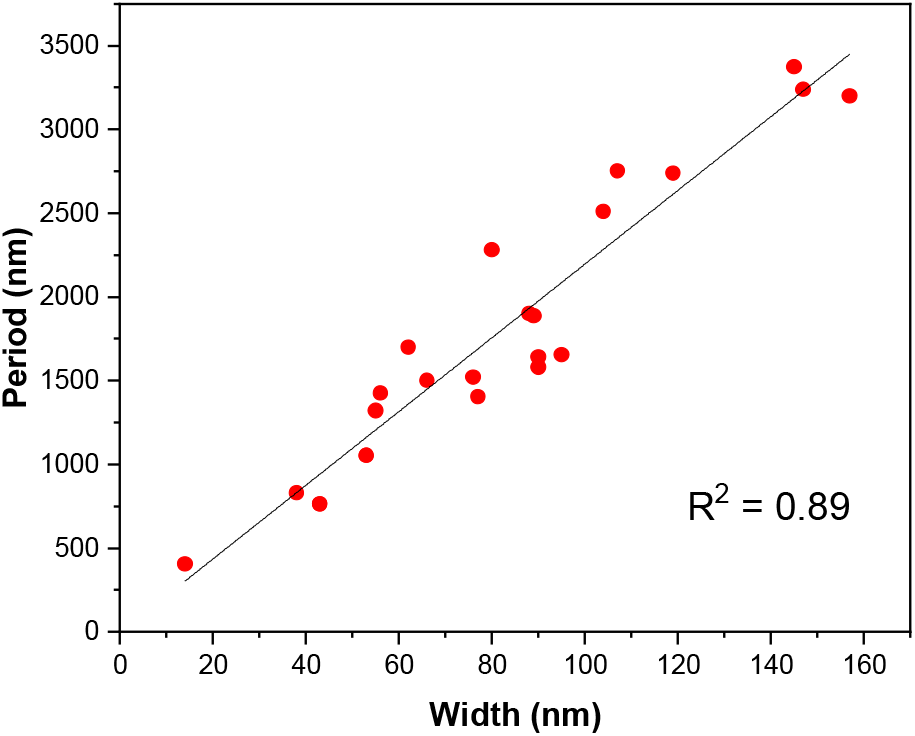
Helical pitch of twisted ribbons plotted against width of the fibrils. This result shows a linear relation between fibril width and helical pitch of α-syn amyloid fibrils formed in the presence of DMPS vesicles.

The large widths observed for the twisted and helical ribbon polymorphs suggest that they are composed of a number of protofilaments. This observation is consistent with models that consider hierarchical assembly of protofilaments into larger amyloid fibril polymorphs^15,72^. Indeed, we observe several helical and twisted ribbons that appear to split along the length of the fibril (**Figure 6**). Moreover, we see that this splitting can happen at multiple locations along the length of single fibril. The individual strands of these “frayed” fibrils have a larger width than that observed for protofilaments (3 nm). This implies that fibrils consisting of many protofibrils may further assemble to produce giant fibrils in a hierarchical assembly.

**Figure 6.**
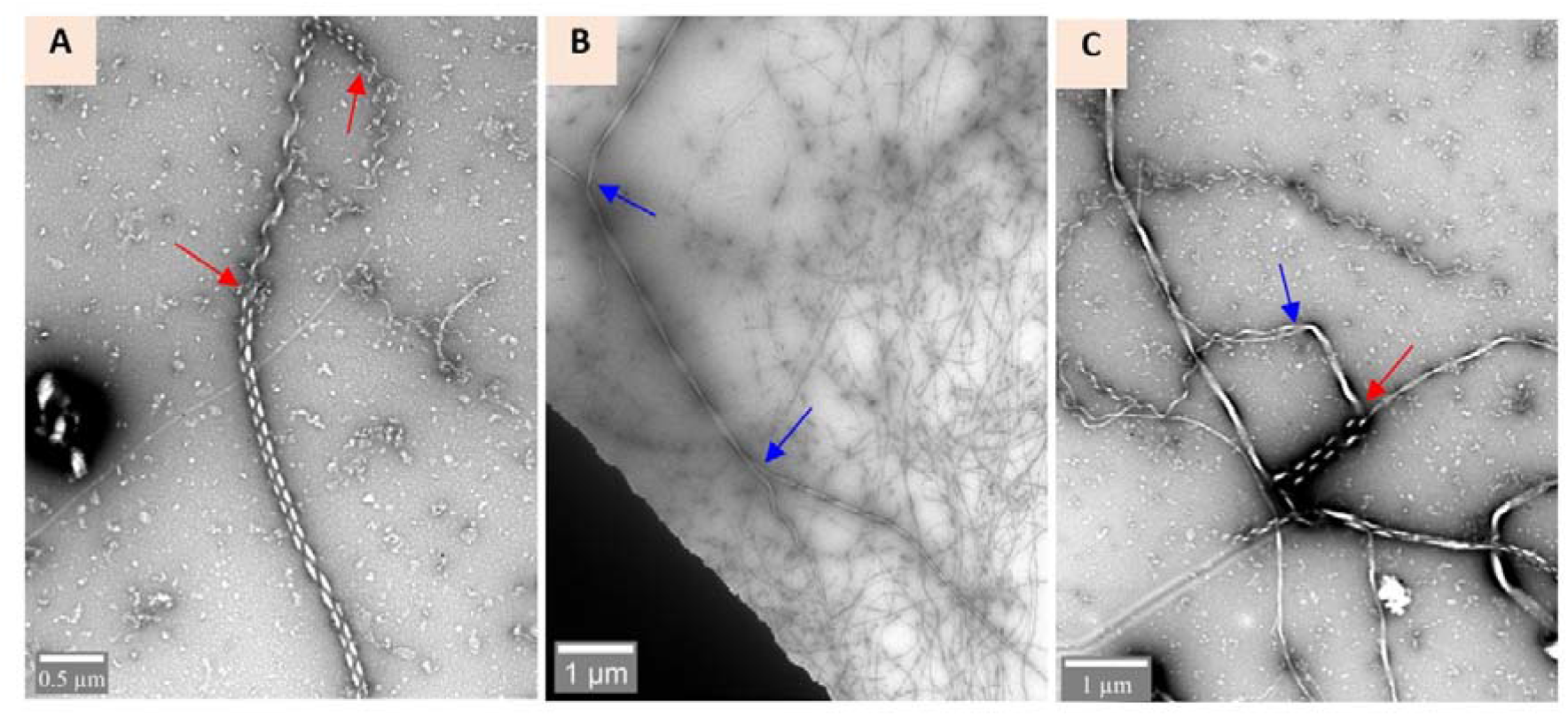
TEM imaging of fibril splitting: Fibrils show splitting along their length as shown in **A-C**. Splitting was observed for both helical (marked with red arrow) and twisted (marked with blue arrow) fibrils. This result shows that observed giant fibrils are composed of mature fibrils.

### Diverse polymorphs form at early stages of aggregation

In order to better understand the structural kinetics of these highly polymorphic structures, we performed time dependent imaging of α-syn aggregating in the presence of DMPS vesicles. We monitored the kinetics of aggregation using ThT fluorescence assays and sampled aliquots from these experiments for TEM analysis after 4 h and 15 h of incubation. TEM image analysis of these samples showed the presence of helical fibrils and twisted ribbons at both time points (**Fig 7A-B; 7C-D**), however, these fibrils were only a few microns in length, significantly shorter than fibrils observed at the plateau phase. It has been shown that twisted ribbons are metastable and tend to form helical fibrils which are thermodynamically more stable^64^. However, studies with other amyloid systems show these forms take considerable time to convert from twisted to helical forms^64,65^. In our study we find that these forms coexist with one another and appear at very early times in the aggregation kinetics. This strongly suggests interaction with lipid molecules helps the twisted ribbon to cross the thermodynamic barrier and convert into the more stable helical form.

**Figure 7.**
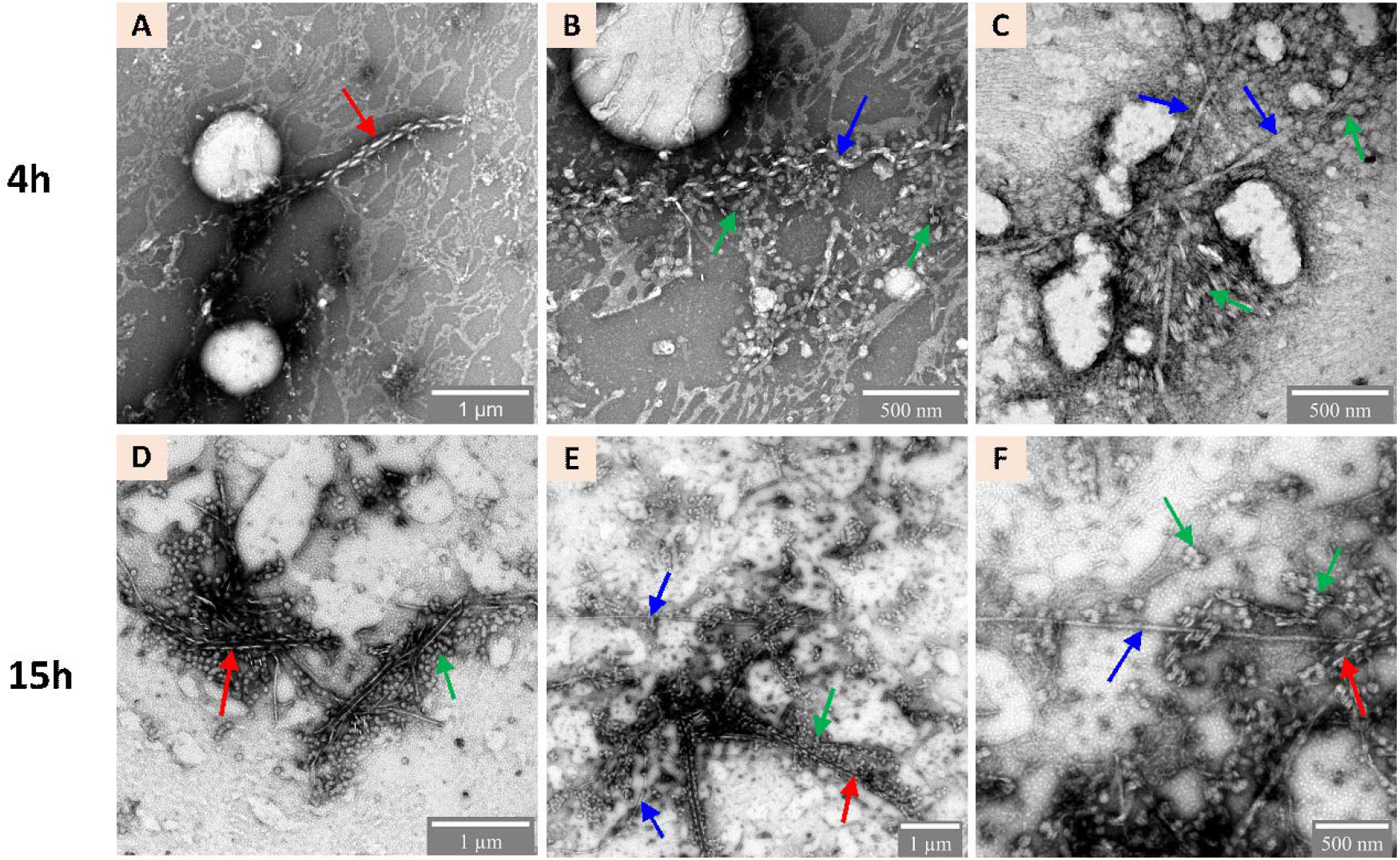
TEM imaging of sample during the early growth phase: TEM analysis of samples after 4h (A-C) and 15h (D-F) of incubation shows presence of helical (red arrow) and twisted ribbons (blue arrow) embedded in nanodiscs-like structures (green arrow).

Interestingly, in addition to the structures already discussed, we also observed the presence of nanodisc-like structures which were closely associated with most fibrils. The analysis of these discs reveal that they possess an internal spacing of 3-4 nm (**Fig. S5**). Such nanodisc structures are known to form in the presence of phospholipids and membrane scaffolding proteins (MSPs)^74^. A typical example of MSP is apolipoprotein, which shares significant sequence homology with α-syn^75^ and a common ability to induce membrane curvature^76^. It has been also shown that α-syn can form such structures in the presence of some phospholipids^74,77,78^. In our experiment the absence of these discoid structures in the late aggregation phase suggests these structures are transient in nature.

### Fibril formed in the presence of DMPS vesicles co-localize with lipid molecules

Previous *in vitro* studies have shown that co-assembly can occur between lipid molecules and α-syn during amyloid fibril formation^43,79^, and in a recent study it has been shown that Lewy bodies, the hallmark of Parkinson’s disease, consist of highly crowded mixtures of organellar and membranous features that contain inclusions of α-syn^80^. Interestingly, in another study, incubation of α-syn on a supported bilayer of phospholipid membrane resulted in the aggregation of α-syn accompanied by lipid extraction from the membrane, causing membrane disruption and clustering of lipid molecules around growing α-syn aggregates^41^. To see whether α-syn fibrils can incorporate lipid, we employed total internal reflection fluorescence microscopy (TIRFM) to image individual α-syn aggregates (Single Aggregate Visualization by Enhancement (SAVE) imaging)^81^. For these experiments, we utilised DMPS liposomes that had been conjugated with biotinylated lipid. These biotinylated liposomes were added to our α-syn samples and allowed to undergo fibrilization. Samples were collected and incubated with Alexa Fluor 647 labelled streptavidin (1 nM) and 5 uM ThT to allow for simultaneous imaging and localization of lipids and fibrils, respectively. We found a strong co-localiztion of the lipids and fibrils as shown in Figs. **8A** and **8B**. In contrast, when fibrils were prepared without biotinylated lipids, no co-localization is observed (**Figs. S6**). The association quotient, *Q*^82–84^ (see single-molecule data analysis in Methods), is a measure of the level of coincidence of the fibrils and lipids. For these samples, *Q* = 0.43 ± 0.17 (mean ± S.D., *n* = 27 images), indicating that 43% of the fibrils identified contained biotinylated lipid. For the control in which biotinylated lipid was not included, *Q =* −0.02 ± 0.04 (mean ± S.D., *n* = 5 images).

**Figure 8.**
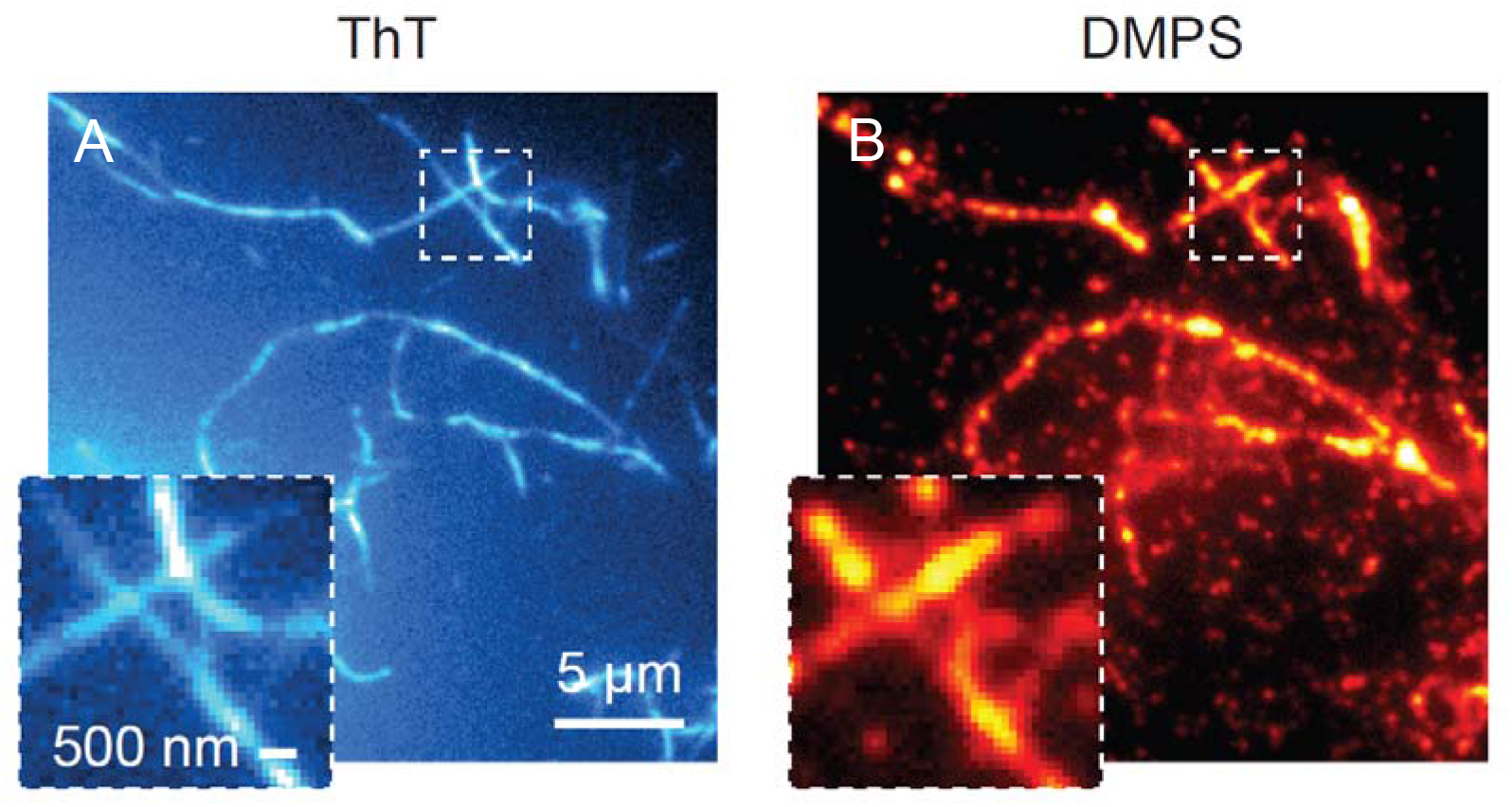
α-Syn amyloid fibrils association with lipid observed by TIRFM imaging. 0.5 μM α-syn amyloid fibrils and in the presence of 4 μM DMPS vesicles (containing 2 mol % of biotinylated lipid) in the presence of 1 nM streptavidin-AF647 and 5μM ThT **(A)**Images were recorded for 50 frames from the red channel (AF647 emission) with 641 nm illumination, followed by **(B)**green channel (ThT emission) with 405 nm illumination.

Taken together, these imaging techniques demonstrate that in the presence of DMPS lipid vesicles, α-syn forms aggregates with nanodisc-like structures and that the growing fibrils can directly incorporate lipid molecules. Further study of these highly polymorphic α-syn amyloid fibril variants can better illuminate the processes that govern lipid-induced conversion of α-syn into potentially pathological forms.

## Conclusions

With a growing number of studies, it is becoming increasingly clear that heterogenous nucleation plays an important role in amyloid formation in different amyloid forming proteins. Most importantly, many studies have shown interactions between lipid bilayers and amyloid forming proteins is a common triggering event in the formation of amyloid fibrils and other cytotoxic species^85–87^. Current evidence suggests that lipid membranes provide a platform where the binding of amyloidogenic protein increases the local concentration of bound protein molecules and thus facilitates the formation of toxic aggregates. In the work reported here, we have explored the effect of physiological relevant lipid DMPS on α-syn fibrillization.

Analysis of ThT fluorescence data showed that with an increase in the lipid/protein ratio, lagtime was significantly decreased. However, the presence of lipid does not affect the variability of the lag-phase (**Fig. 1A**). TEM imaging of the samples revealed the presence of a large spectrum of amyloid fibril structures when α-syn was aggregated in the co-presence of DMPS vesicles. We identified multiple structures such as thin and curly fibrils, twisted and helical ribbons, nanotubes and flat sheets (**Fig 2D - 2I**). Although all of these structures have been reported before for amyloid forming proteins and short peptides, no study has described the presence of so many variants under identical environmental conditions. When we carried out the reaction in the absence of DMPS, we observed either small oligomeric aggregates or straight fibrils with rod-like morphologies (**Fig. 2A-Fig. 2C**). Quantitative analysis of these fibrils showed they possesses a wide spectrum of lengths and widths as compared to fibrils in the absence of DMPS (**Fig. 4**). Further analysis of the width-to-period relation of twisted ribbons showed a linear relation, which suggest these fibrils have a different number of constituent protofilaments (**Fig 5**). These results clearly show that the presence of DMPS vesicles not only modulates the kinetics of fibril formation of α-syn but importantly, its interaction leads to the formation of polymorphic fibril forms. Polymorphic forms of α-syn are known to be present under *in vivo* conditions and they have been linked to different pathological conditions^22,25,88–90^, however, factors responsible for formation of these polymorphs under *in vivo* conditions are not known. The findings of our present study highlight the possible role lipids play in the formation of α-syn polymorphs under *in vivo* conditions.

Another principal finding of this work is the observation of the presence of nano-disc like structures at the early stage of aggregation kinetics (**Fig 7 and S3**). Our observation that that these discoid structures disappear in the plateau phase of the kinetics as well as previous reports of lipid molecule clustering on growing α-syn fibrils^41^ prompted us to investigate if lipid molecules are being directly incorporated into amyloid fibrils. Experiments carried out with TIRF microscopy shows a high-level of co-localization of lipid and amyloid fibrils. Thus, we confirmed that lipids can be incorporated into the structure of amyloid fibrils during their growth process.

In summary, we have shown that under *in vitro* conditions, lipid-protein interactions play an important role in the formation of polymorphs of α-syn amyloid fibrils. Further study is needed to understand the mechanism of the lipid-induced conversion of α-syn into its polymorphic forms. The knowledge obtained from such a study is also expected to provide crucial information on the widely observed lipid-induced conversion of amyloid forming proteins into their toxic forms.

## Methods

### Reagents

1,2-dimyristoyl-sn-glycero-3-phospho-L-serine (DMPS) lyophilized powder and 1,2-distearoyl-sn-glycero-3-phosphoethanol-amine-N-[amino(polyethyleneglycol0)] (DSPE-PEG) were purchased from Avanti Polar Lipid, Inc. Streptavidin, Alexa Fluor 647 conjugate was purchased from ThermoFisher Scientific. Tris base, ethylenediaminetetraacetic acid (EDTA), Thioflavin T, Chloroform, Methanol, and Hydrochloric acid (HCl) and other chemicals were purchased from Sigma Aldrich.

### Protein purification

Recombinant WT α-syn was expressed in *Escherichia Coli* and purified as described previously^91^.

### Liposome preparation

About 1 mg DMPS were obtained in a glass test-tube and dissolved in chloroform:methanol (3:1 v/v) solvent. Thin layer of lipid was obtained by constant rotating of test-tube, while gentle stream of nitrogen gas was used to evaporate the solvent. To remove the any trace amount of solvent, test-tube was placed in the vacuum for minimum of 4h. Liposome biotinylation was achieved by adding 2 mol % of biotinylated lipid (DSPE-PEG Biotin, Avanti) to DMPS lipid. The thin lipid layer was then hydrated with appropriate amount of 25 mM tris buffer, pH 7.4 to make a stock concentration of 2 mM and vortexed for 2 mins to dissolve the lipid layer in the buffer. Sample was frozen and thawed for 4 cycle on dry ice and water-bath at 40 °C. Small unilamellar vesicles (SUVs) were prepared by sonication (VWR ultrasonic cleaner) for 30 minutes. Size of the liposomes was characterized by dynamic light scattering and were shown to consist of small and large peak centered at a diameter of 17 nm (small peak) and 109 nm (large peak), which is similar to previous report^32^ (**Fig. S1**).

### Dynamic light scattering (DLS)

Size distribution of lipid vesicles was determined by an ALV/CGS-3 platform-based goniometer system (ALV-GmbH). This instrument was equipped with a 22 mW HeNe laser with a wavelength of 633 nm, and backscattered light was detected at an angle of 90° at room temperature. DMPS stock solutions were diluted to 60 μM in filtered 25 mM phosphate buffer (pH 7.4), transferred into glass test tube and placed in the measurement cell. Three measurements were performed for the sample for 300 seconds at room temperature and average values were used for analysing the data.

### Aggregation assay

ThT fluorescence time-course measurements were performed in 96-well microliter plates using Fluostar Omega microplate reader (BMG Labtech) with excitation and emission wavelengths set to 450 nm and 485 nm. 50 μM WT α-syn samples were prepared in the absence and presence of 25 μM, 50 μM, 100 μM, 200 μM and 400 μM of DMPS vesicles with 25 μM ThT in 25 mM Tris buffer (pH 7.4). Each sample-well contained 100 μL of the reaction mixture and spontaneous aggregation was induced by incubation at 30 °C without shaking.

### Transmission electron microscopy

At the end of aggregation assay, aggregated sample was diluted to 1 μM concentration. An aliquot (7 μL) of diluted sample was placed onto carbon support film 300 Mesh 3 mm copper grids (TAAB) for 3 min and blot dried. The TEM grids were subsequently stained using 7 μL of 1 % uranyl acetate for 2 min followed by blot drying. Samples were imaged on a Thermo Fisher Scientific Tecnai F20 electron microscope (200 kV, field emission gun) equipped with an 8k x 8k CMOS camera (TVIPS F816).

### CD Spectroscopy

A Jasco J-810 spectrometer was used for CD measurements. Samples containing 10 μM WT α-syn in presence of different amount of DMPS (0, 10, 20, 30, 40, 50, 60, 70, 80, 90, 100, 110, 120, 130, 140, 150, 160, 170, 200, 250 and 300 μM) were prepared in 25 mM phosphate buffer (pH 7.4). Experiments were performed at 25 °C using a quartz cuvette with 1 mm path length at a scan rate 20 nm min^−1^ with a data pitch of 0.1 nm and a digital integration time of 1s. A total five scan were accumulated and averaged for a final data.

### Slide preparation for single-molecule analysis

Glass coverslips (22 × 22 mm, VWR international, USA) were cleaned using an argon plasma cleaner (Zepto, Diener, Germany) for 40 minutes to remove any fluorescent residue. Frame-seal slide chambers (9 x 9 cm^2^, Bio-Rad, Hercules, USA) were affixed to the glass, 50 μl of poly-L-lysine (70-150k molecular weight, Sigma-Aldrich) was added and incubated at room temperature for at least 15 minutes. The coverslips were washed twice with 20 mM filtered buffer (20 mM Tris, 100 mM NaCl, pH 7.4) immediate before adding the sample for imaging.

### Total internal reflection fluorescence microscopy

Single-molecule imaging was performed using a custom-built single-molecule TIRF microscope, which restricts the illumination to within ~200 nm of the sample slide. The fluorophores were excited at either 405 nm (ThT), or 638 nm (AF647). Collimated laser light at wavelengths of 405 nm (Cobolt MLD Series 405-250 Diode Laser System, Cobolt AB, Solna, Sweden), and 638 nm (Cobolt MLD Series 638-140 Diode Laser System, Cobolt AB, Solna, Sweden) were aligned and directed parallel to the optical axis at the edge of a 1.49 NA TIRF objective (CFI Apochromat TIRF 60XC Oil, Nikon, Japan), mounted on an inverted Nikon TI2 microscope (Nikon, Japan). The microscope was fitted with a perfect focus system to autocorrect the z-stage drift during imaging. Fluorescence collected by the same objective was separated from the returning TIR beam by a dichroic mirror (Di01-R405/488/ 561/635 (Semrock, Rochester, NY, USA), and was passed through appropriate filters (405 nm: BLP01-488R-25 (Semrock, Rochester, NY, USA), 638 nm: FF01-692/40-25 (Semrock, Rochester, NY, USA). The fluorescence was then passed through a 2.5× beam expander and recorded on an EMCCD camera (Delta Evolve 512, Photometrics, Tucson, AZ, USA) operating in frame transfer mode (EMGain = 11.5 e-/ ADU and 250 ADU/photon). Each pixel was 103 nm in length. Images were recorded over 10 frames with an exposure time of 50 ms with 638 nm (~50 W cm^−2^) illumination, followed by 405 nm excitation (~100 W cm^−2^). The microscope was automated using the open source microscopy platform Micromanager.

Sample collected from a 96 well-plate after completion of aggregation reaction was diluted 100 × and incubated for 10 minutes in presence of 1 nM Straptavidin-AF647 (ThermoFisher Scientific). It was then centrifuged at 14k rpm for 10 minutes and the pellet was redissolved in 5 μM ThT and placed on glass coverslip for imaging.

### Single-molecule data analysis

The data were analysed using a custom-written code in Igor Pro (Wavemetrics). Each tiff image stack was first averaged over the 10 frames in each channel, before having the background subtracted. A manual threshold value of 1000 ADU (ThT channel) was used to distinguish fibrils from background. A clustering algorithm (Density-based spatial clustering of applications with noise^92^) was used to identify individual fibrils. For each fibril, the corresponding fluorescence was analysed in the AF647 channel for each pixel, and a positive coincidence was defined if at least one of these had a value greater than an applied threshold of 1500 ADU in the AF647 channel. To account for chance coincidence (i.e. due to the fibrils and AF647-tagged liposomes being in close proximity), the same analysis was performed with the AF647 images being rotated 90° and translated 100 pixels. For each image set, the association quotient, Q^82–84^, which is a measure of coincidence, was calculated according to Equation 1.

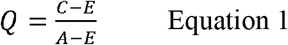

Where C is the number of coincident events, E the number of chance events, and A the total number of fibrils detected.

## Acknowledgments

This work was supported by The Royal Society UK, and Science and Engineering Research Board (SERB), India. M.H.H. acknowledges funding from the UK Dementia Research Institute and UCB to build the total internal reflection fluorescence microscope (https://www.ucb.com).

## References

1. Graumann, P. L. Cytoskeletal elements in bacteria. Curr. Opin. Microbiol. 7, 565–571 (2004).

2. Kueh, H. Y. & Mitchison, T. J. Structural Plasticity in Actin and Tubulin Polymer Dynamics. Science (80-.). 325, 960–963 (2009).

3. Ryan, E. A., Mockros, L. F., Weisel, J. W. & Lorand, L. Structural Origins of Clot Rheology. Biophys. J. 77, 2813–2826 (1999).

4. Buehler, M. J. Nature designs tough collagen: Explaining the nanostructure of collagen fibrils. Proc. Natl. Acad. Sci. U. S. A. 103, 12285–12290 (2006).

5. Römer, L. & Scheibel, T. The elaborate structure of spider silk: structure and function of a natural high performance fiber. Prion 2, 154–161 (2008).

6. Chiti, F. & Dobson, C. M. Protein Misfolding, Functional Amyloid, and Human Disease. Annu. Rev. Biochem. 75, 333–366 (2006).

7. Knowles, T. P. J. & Mezzenga, R. Amyloid Fibrils as Building Blocks for Natural and Artificial Functional Materials. Adv. Mater. 28, 6546–6561 (2016).

8. Knowles, T. P. J. & Buehler, M. J. Nanomechanics of functional and pathological amyloid materials. Nat. Nanotechnol. 6, 469–479 (2011).

9. Khurana, R. et al. A general model for amyloid fibril assembly based on morphological studies using atomic force microscopy. Biophys. J. 85, 1135–44 (2003).

10. Knowles, T. P. J., Vendruscolo, M. & Dobson, C. M. The amyloid state and its association with protein misfolding diseases. Nat. Rev. Mol. Cell Biol. 15, 384–396 (2014).

11. Eisenberg, D. & Jucker, M. The amyloid state of proteins in human diseases. Cell 148, 1188–1203 (2012).

12. Nelson, R. et al. Structure of the cross-β spine of amyloid-like fibrils. Nature 435, 773–778 (2005).

13. Li, B. et al. Cryo-EM of full-length α-synuclein reveals fibril polymorphs with a common structural kernel. Nat. Commun. 9, 1–10 (2018).

14. Elizabeth L. Guenther, Peng Ge, Hamilton Trinh, Michael. Sawaya, Duilio Cascio, David R. Boyer, Tamir Gonen, Z. Hong Zhou, and D. S. E. Atomic-level evidence for packing and positional amyloid polymorphism by segment from TDP-43 RRM2. Nat Struct Mol Biol. 25, 311–319 (2018).

15. Fitzpatricka, A. W. P. et al. Atomic structure and hierarchical assembly of a cross-β amyloid fibril. Proc Natl Acad Sci USA 5590, 5468–5473 (2013).

16. Anthony W.P., et al. Cryo-EM structures of Tau filaments from Alzheimer’s disease brain. Nature 547, 185–190 (2017).

17. Chien, P., Weissman, J. S. & DePace, A. H. Emerging Principles of Conformation-Based Prion Inheritance. Annu. Rev. Biochem. 73, 617–656 (2004).

18. Qiang, W., Yau, W. M., Lu, J. X., Collinge, J. & Tycko, R. Structural variation in amyloid-β fibrils from Alzheimer’s disease clinical subtypes. Nature 541, 217–221 (2017).

19. Adamcik, J. & Mezzenga, R. Amyloid Polymorphism in the Protein Folding and Aggregation Energy Landscape. Angew. Chemie - Int. Ed. 57, 8370–8382 (2018).

20. Lara, C., Adamcik, J., Jordens, S. & Mezzenga, R. General self-assembly mechanism converting hydrolyzed globular proteins into giant multistranded amyloid ribbons. Biomacromolecules 12, 1868–1875 (2011).

21. Marcus Fändrich, Sofie Nyström, K. Peter. Nilsson, Anja Böckmann, Harry LeVine III, and P. H. Amyloid fibril polymorphism - a challenge for molecular imaging and therapy. J Intern Med. 283, 218–237 (2018).

22. Lau, A. et al. α-Synuclein strains target distinct brain regions and cell types. Nat. Neurosci. 23, 21–31 (2020).

23. Lu, J. X. et al. Molecular structure of ß-amyloid fibrils in alzheimer’s disease brain tissue. Cell 154, 1257 (2013).

24. Aneta T. Petkova, Richard D. Leapman, Zhihong Guo, Wai-Ming Yau, Mark P. Mattson, R. T. Self-Propagating, Molecular-Level Polymorphism in Alzheimer’s □-Amyloid Fibrils. Science (80-.). 307, 262–265 (2005).

25. Peelaerts, W. & Baekelandt, V. □-Synuclein strains and the variable pathologies of synucleinopathies. J. Neurochem. 139, 256–274 (2016).

26. Goedert, M., Jakes, R. & Spillantini, M. G. The Synucleinopathies: Twenty Years on. J. Parkinsons. Dis. 7, S53–S71 (2017).

27. Galvagnion, C. The Role of Lipids Interacting with α-Synuclein in the Pathogenesis of Parkinson’s Disease. J. Parkinsons. Dis. 7, 433–450 (2017).

28. Bendor, J. T., Logan, T. P. & Edwards, R. H. The function of α-synuclein. Neuron 79, 1044–1066 (2013).

29. Beatman, E. L. et al. Alpha-Synuclein Expression Restricts RNA Viral Infections in the Brain. J. Virol. 90, 2767–2782 (2016).

30. de Diego, I., Peleg, S. & Fuchs, B. The role of lipids in aging-related metabolic changes. Chem. Phys. Lipids 222, 59–69 (2019).

31. Shulman, J. M., De Jager, P. L. & Feany, M. B. Parkinson’s disease: Genetics and pathogenesis. Annu. Rev. Pathol. Mech. Dis. 6, 193–222 (2011).

32. Galvagnion, C. et al. Lipid vesicles trigger α-synuclein aggregation by stimulating primary nucleation. Nat. Chem. Biol. 11, 229–234 (2015).

33. Ross, B. M., Mamalias, N., Moszczynska, A., Rajput, A. H. & Kish, S. J. Elevated activity of phospholipid biosynthetic enzymes in substantia nigra of patients with Parkinson’s disease. Neuroscience 102, 899–904 (2001).

34. Eugène, S., Xue, W. F., Robert, P. & Doumic, M. Insights into the variability of nucleated amyloid polymerization by a minimalistic model of stochastic protein assembly. J Chem Phys 144, doi: 10.1063/1.4947472 (2016).

35. Gaspar, R., Pallbo, J., Weininger, U., Linse, S. & Sparr, E. Ganglioside lipids accelerate α-synuclein amyloid formation. Biochim. Biophys. Acta - Proteins Proteomics 1866, 1062–1072 (2018).

36. Benjamin G. Wilhelm, Sunit Mandad, Sven Truckenbrodt, Katharina Kröhnert, Christina Schäfer, Burkhard Rammner, Seong Joo Koo, Gala A. Claßen, Michael Krauss, Volker Haucke, Henning Urlaub, S. O. R. Composition of isolated synaptic boutons reveals the amounts of vesicle trafficking proteins. Science (80-.). 344, 1023–1028 (2014).

37. Wei, G. et al. Self-assembling peptide and protein amyloids: From structure to tailored function in nanotechnology. Chem. Soc. Rev. 46, 4661–4708 (2017).

38. Usov, I., Adamcik, J. & Mezzenga, R. Polymorphism complexity and handedness inversion in serum albumin amyloid fibrils. ACS Nano 7, 10465–10474 (2013).

39. Adamcik, J. & Mezzenga, R. Proteins Fibrils from a Polymer Physics Perspective. Macromolecules 45, 1137–1150 (2012).

40. Adamcik, J. et al. Microtubule-Binding R3 Fragment from Tau Self-Assembles into Giant Multistranded Amyloid Ribbons. Angew. Chemie - Int. Ed. 55, 618–622 (2016).

41. Reynolds, N. P. et al. Mechanism of membrane interaction and disruption by α-synuclein. J. Am. Chem. Soc. 133, 19366–19375 (2011).

42. Jensen, P. H., Nielsen, M. S., Jakes, R., Dotti, C. G. & Goedert, M. Binding of α-synuclein to brain vesicles is abolished by familial Parkinson’s disease mutation. J. Biol. Chem. 273, 26292–26294 (1998).

43. Galvagnion, C. et al. Lipid Dynamics and Phase Transition within α-Synuclein Amyloid Fibrils. J. Phys. Chem. Lett. 10, 7872–7877 (2019).

44. Martinez, Z., Zhu, M., Han, S. & Fink, A. L. GM1 specifically interacts with α-synuclein and inhibits fibrillation. Biochemistry 46, 1868–1877 (2007).

45. Iyer, A. & Claessens, M. M. A. E. Disruptive membrane interactions of alpha-synuclein aggregates. Biochim. Biophys. Acta - Proteins Proteomics 1867, 468–482 (2019).

46. Grey, M., Linse, S., Nilsson, H., Brundin, P. & Sparr, E. Membrane interaction of α-synuclein in different aggregation states. J. Parkinsons. Dis. 1, 359–371 (2011).

47. Ugalde, C. L., Lawson, V. A., Finkelstein, D. I. & Hill, A. F. The role of lipids in α-synuclein misfolding and neurotoxicity. J. Biol. Chem. 294, 9016–9028 (2019).

48. Ryan, T. et al. Cardiolipin exposure on the outer mitochondrial membrane modulates α-synuclein. Nat. Commun. 9, 1–17 (2018).

49. Galvagnion, C. et al. Chemical properties of lipids strongly affect the kinetics of the membrane-induced aggregation of α-synuclein. Proc. Natl. Acad. Sci. 113, 7065–7070 (2016).

50. Jo, E., McLaurin, J. A., Yip, C. M., St. George-Hyslop, P. & Fraser, P. E. α-Synuclein membrane interactions and lipid specificity. J. Biol. Chem. 275, 34328–34334 (2000).

51. Pirc, K. & Ulrih, N. P. α-Synuclein interactions with phospholipid model membranes: Key roles for electrostatic interactions and lipid-bilayer structure. Biochim. Biophys. Acta - Biomembr. 1848, 2002–2012 (2015).

52. Zhu, M. & Fink, A. L. Lipid binding inhibits α-synuclein fibril formation. J. Biol. Chem. 278, 16873–16877 (2003).

53. Meade, R. M., Williams, R. J. & Mason, J. M. A series of helical α-synuclein fibril polymorphs are populated in the presence of lipid vesicles. npj Park. Dis. 6, (2020).

54. Guerrero-Ferreira, R. et al. Two new polymorphic structures of human full-length alpha-synuclein fibrils solved by cryo-electron microscopy. Elife 8, 1–24 (2019).

55. Periole, X. et al. Energetics Underlying Twist Polymorphisms in Amyloid Fibrils. J. Phys. Chem. B 122, 1081–1091 (2018).

56. Dobson, C. M. Protein folding and misfloding. Nature 426, 884–890 (2003).

57. Fusco, G. et al. Direct observation of the three regions in α-synuclein that determine its membrane-bound behaviour. Nat. Commun. 5, 1–17 (2014).

58. Ferreon, A. C. M. & Deniz, A. A. α-Synuclein multistate folding thermodynamics: Implications for protein misfolding and aggregation. Biochemistry 46, 4499–4509 (2007).

59. Anderson, V. L., Ramlall, T. F., Rospigliosi, C. C., Webb, W. W. & Eliezer, D. Identification of a helical intermediate in trifluoroethanol-induced alpha-synuclein aggregation. Proc. Natl. Acad. Sci. 107, 18850–18855 (2010).

60. Munishkina, L. A., Phelan, C., Uversky, V. N. & Fink, A. L. Conformational behavior and aggregation of α-synuclein in organic solvents: Modeling the effects of membranes. Biochemistry 42, 2720–2730 (2003).

61. Lu, K., Jacob, J., Thiyagarajan, P., Conticello, V. P. & Lynn, D. G. Exploiting amyloid fibril lamination for nanotube self-assembly. J. Am. Chem. Soc. 125, 6391–6393 (2003).

62. Aggeli, A. et al. Hierarchical self-assembly of chiral rod-like molecules as a model for peptide-sheet tapes, ribbons, fibrils, and fibers. Proc. Natl. Acad. Sci. 98, 11857–11862 (2001).

63. Lamm, M. S., Rajagopal, K., Schneider, J. P. & Pochan, D. J. Laminated morphology of nontwisting β-sheet fibrils constructed via peptide self-assembly. J. Am. Chem. Soc. 127, 16692–16700 (2005).

64. Pashuck, E. T. & Stupp, S. I. Direct observation of morphological tranformation from twisted ribbons into helical ribbons. J. Am. Chem. Soc. 132, 8819–8821 (2010).

65. Zhang, S. et al. Coexistence of ribbon and helical fibrils originating from hIAPP 20-29 revealed by quantitative nanomechanical atomic force microscopy. Proc. Natl. Acad. Sci. U. S. A. 110, 2798–2803 (2013).

66. Serpell, L. C., Berriman, J., Jakes, R., Goedert, M. & Crowther, R. A. Fiber diffraction of synthetic alpha-synuclein filaments shows amyloid-like cross-beta conformation. Proc. Natl. Acad. Sci. 97, 4897–4902 (2000).

67. Paravastu, A. K., Leapman, R. D., Yau, W.-M. & Tycko, R. Molecular structural basis for polymorphism in Alzheimer’s β-amyloid fibrils. Proc. Natl. Acad. Sci. 105, 18349–18354 (2008).

68. Marcus D. Tuttle, et al. Solid-State NMR Structure of a Pathogenic Fibril of Full-Length Human α-Synuclein. Nat Struct Mol Biol. 23, 409–415 (2016).

69. Jose A. Rodriguez, Magdalena I. Ivanova, Michael R. Sawaya, Duilio Cascio, Francis Reyes, Dan Shi, Smriti Sangwan, Elizabeth L. Guenther, Lisa M. Johnson, Meng Zhang, Lin Jiang, Mark A. Arbing, Brent Nannega, Johan Hattne, Julian Whitelegge, Aaron S. Brew, and D. E. Structure of the toxic core of α-synuclein from invisible crystals. Nature 525, 486–490 (2015).

70. Fecchio, C., Palazzi, L. & Polverino de Laureto, P. α-Synuclein and polyunsaturated fatty acids: Molecular basis of the interaction and implication in neurodegeneration. Molecules 23, (2018).

71. Guerrero-Ferreira, R. et al. Cryo-EM structure of alpha-synuclein fibrils. Elife 7, 1–18 (2018).

72. Adamcik, J. et al. Understanding amyloid aggregation by statistical analysis of atomic force microscopy images. Nat. Nanotechnol. 5, 423–428 (2010).

73. Tuttle, M. D. & Comellas, et al. Solid-state NMR structure of a pathogenic fibril of full-length human α-synuclein. Nat. Struct. Mol. Biol. 23, 409–415 (2016).

74. Sligar, I. G. D. and S. G. NANODISCS IN MEMBRANE BIOCHEMISTRY AND BIOPHYSICS. Chem Rev. 176, 4669–4713 (2017).

75. Eichmann, C. et al. Preparation and Characterization of Stable α-Synuclein Lipoprotein Particles. J. Biol. Chem. 291, 8516–8527 (2016).

76. Varkey, J. et al. Membrane Curvature Induction and Tubulation Are Common Features of Synucleins and Apolipoproteins. J. Biol. Chem. 285, 32486–32493 (2010).

77. Varkey, J. et al. α-Synuclein Oligomers With Broken Helical Conformation Form Lipoprotein Nanoparticles. J. Biol. Chem. 288, 17620–17630 (2013).

78. Eichmann, C. et al. Preparation and characterization of stable α-synuclein lipoprotein particles. J. Biol. Chem. 291, 8516–8527 (2016).

79. Hellstrand, E., Nowacka, A., Topgaard, D., Linse, S. & Sparr, E. Membrane Lipid Co-Aggregation with α-Synuclein Fibrils. PLoS One 8, (2013).

80. Shahmoradian, S. H. et al. Lewy pathology in Parkinson’s disease consists of crowded organelles and lipid membranes. Nat. Neurosci. 22, 1099–1109 (2019).

81. Horrocks, M. H. et al. Single-Molecule Imaging of Individual Amyloid Protein Aggregates in Human Biofluids. ACS Chem. Neurosci. 7, 399–406 (2016).

82. Raynes, J. K., Day, L., Crepin, P., Horrocks, M. H. & Carver, J. A. Coaggregation of κ-Casein and ß-Lactoglobulin Produces Morphologically Distinct Amyloid Fibrils. Small 13, doi:10.1002/smll.201603591 (2017).

83. Benson, S. et al. SCOTfluors: Small, Conjugatable, Orthogonal, and Tunable Fluorophores for In Vivo Imaging of Cell Metabolism. Angew. Chemie-Int. Ed. 58, 6911–6915 (2019).

84. Janeczek, A. A. et al. PEGylated liposomes associate with Wnt3A protein and expand putative stem cells in human bone marrow populations. Nanomedicine 12, 845–863 (2017).

85. Gal, N. et al. Lipid bilayers significantly modulate cross-fibrillation of two distinct amyloidogenic peptides. J. Am. Chem. Soc. 135, 13582–13589 (2013).

86. Hebda, J. A. & Miranker, A. D. The Interplay of Catalysis and Toxicity by Amyloid Intermediates on Lipid Bilayers: Insights from Type II Diabetes. Annu. Rev. Biophys. 38, 125–152 (2009).

87. Khondker, A., Alsop, R. J. & Rheinstädter, M. C. Membrane-accelerated Amyloid-β aggregation and formation of cross-β sheets. Membranes (Basel). 7, (2017).

88. Wiltzius, J. J. W. et al. Molecular mechanisms for protein-encoded inheritance. Nat. Struct. Mol. Biol. 16, 973–978 (2009).

89. Shahnawaz, M. et al. Discriminating α-synuclein strains in Parkinson’s disease and multiple system atrophy. Nature 578, (2020).

90. Annamalai, K. et al. Polymorphism of Amyloid Fibrils in Vivo. Angew. Chemie - Int. Ed. 55, 4822–4825 (2016).

91. Phillips, A. S. et al. Conformational dynamics of α-synuclein: Insights from mass spectrometry. Analyst 140, 3070–3081 (2015).

92. Martin Ester, Hans-Peter Kriegel, Jorrg Sander, X. X. A Density-Based Algorithm for Discovering Clusters in Large Spatial Databases with Noise. Proc. Second Int. Conf. Knowl. Discov. Data Min. KDD-96, 226–231

