## Supplementary material for "Lipid-induced polymorphic amyloid fibrils formation by α-synuclein": SI

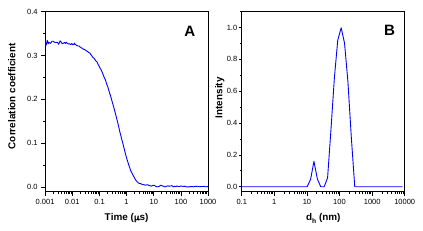

**Figure S1. Dynamic light scattering:** The hydrodynamic diameter (d_h_) of a 60 μM lipid vesicles was determined by dynamic light scattering. **(A)** Correlation function of the sample **(B)** Size distribution of lipid vesicles determined by mass unweighted fitting.

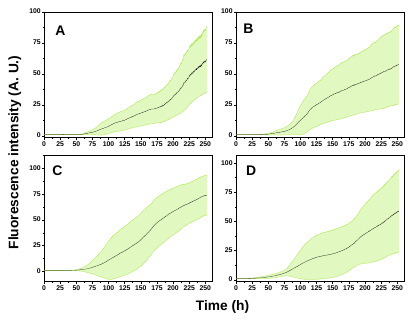

**Figure S2.** Kinetics of fibrillation monitored by the enhancement of thioflavin T fluorescence intensity. Measurements were performed in tris buffer at 30 °C and pH 7.4. Protein concentration was 50 μM in the presence of 25, 50, 100 and 200 μM DMPS vesicles A-D, respectively. Shaded green region is the standard deviation.

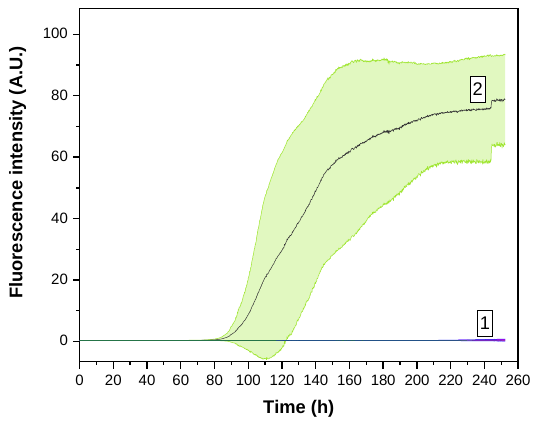

**Figure S3.** Kinetics of α-syn fibrillization in the absence of DMPS vesicles monitored by the enhancement of thioflavin T fluorescence intensity. Measurements were performed at 30 °C and pH 7.4. Curve 1 is mean value when protein concentration was 50 µM and curve 2 is mean value when protein concentration was 100 µM. Shaded green region is the standard deviation.

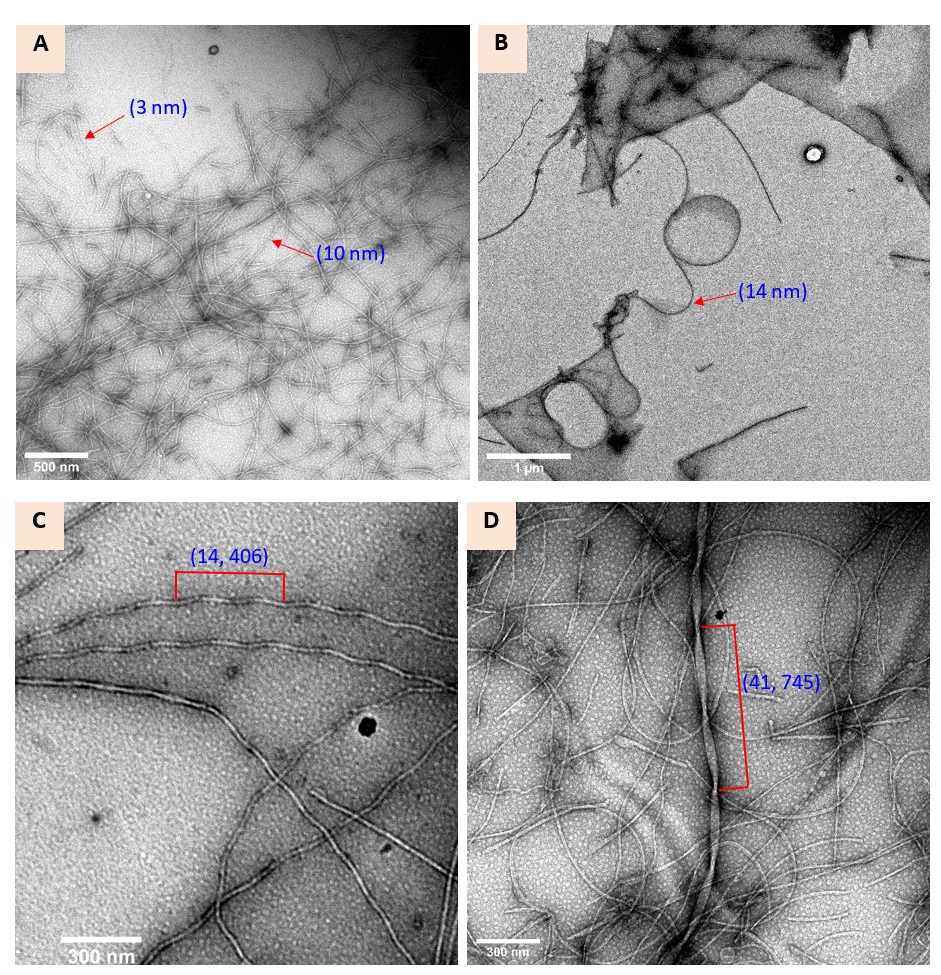

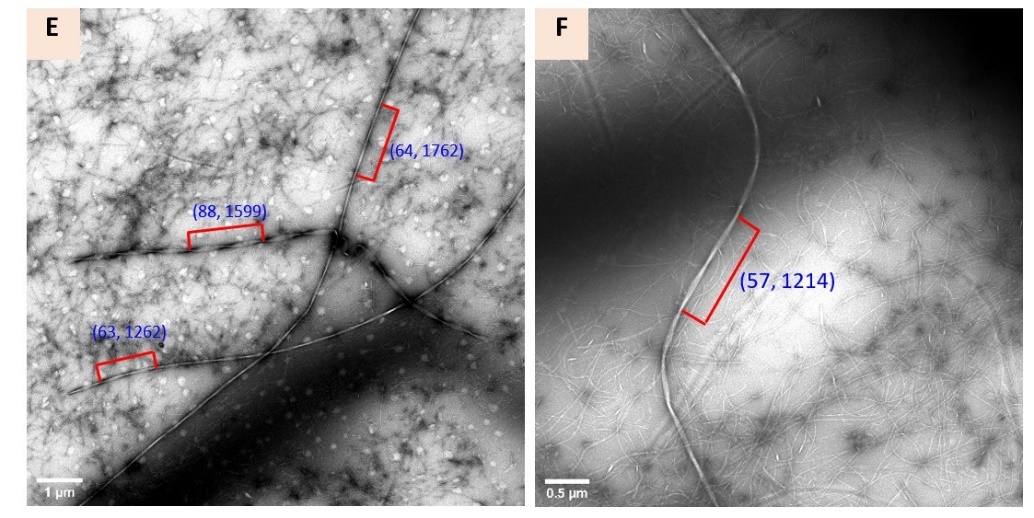

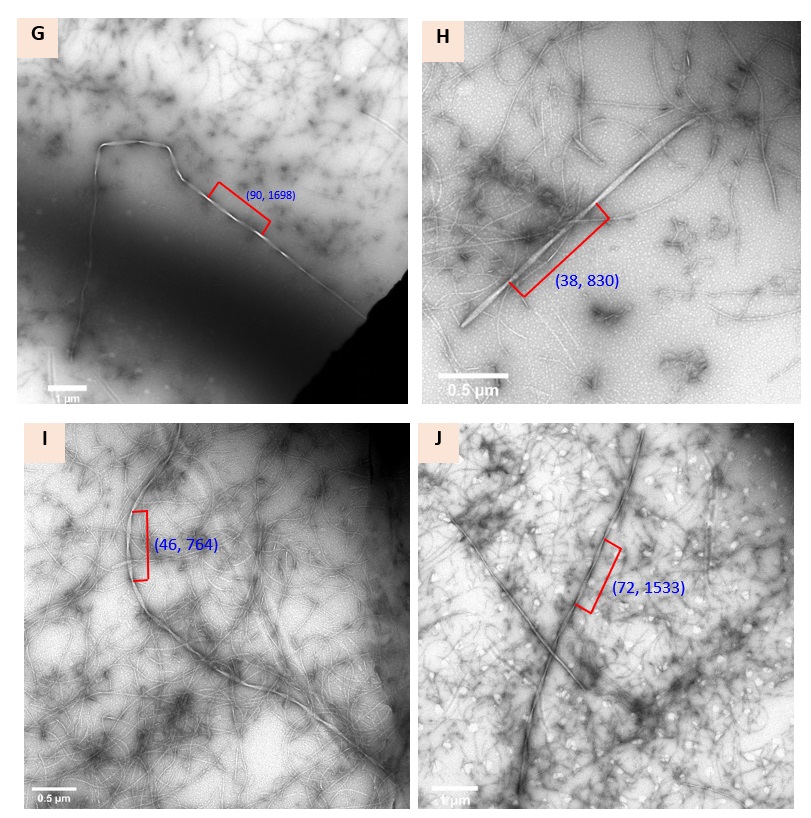

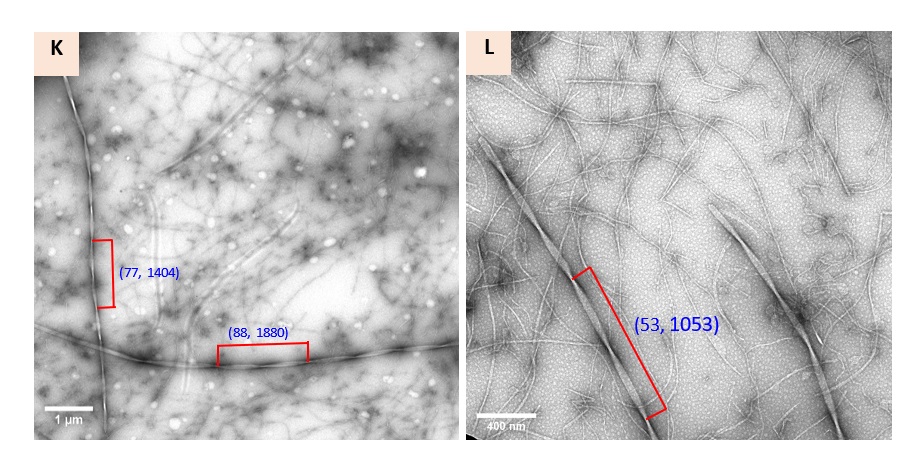

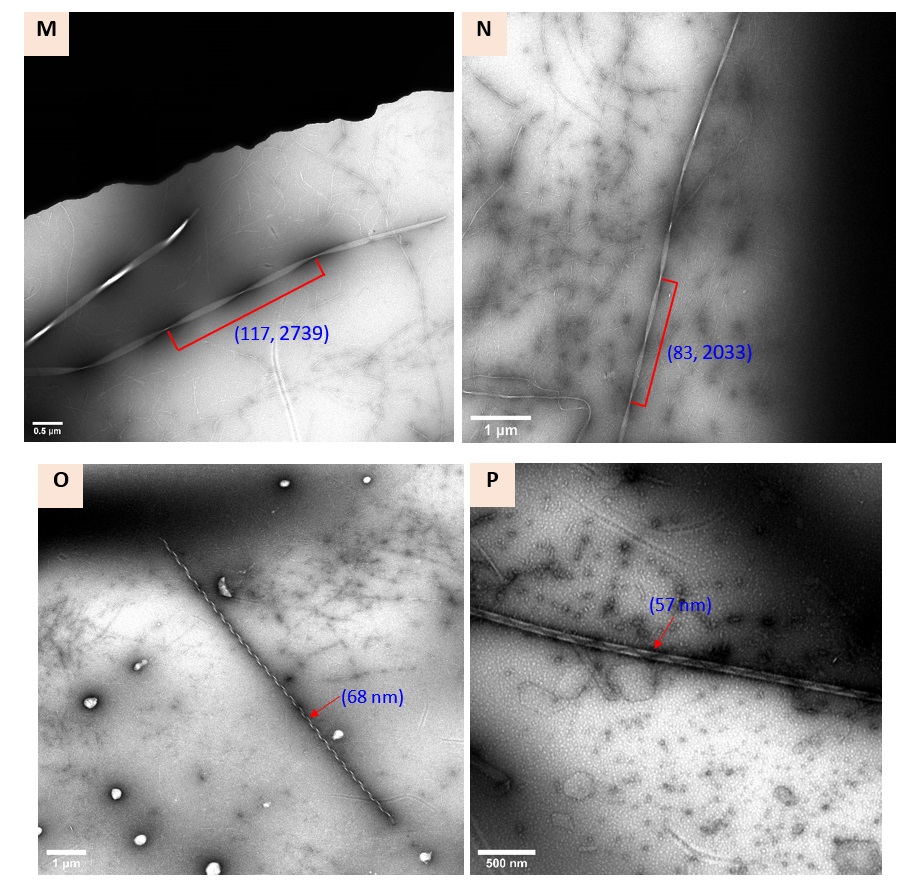

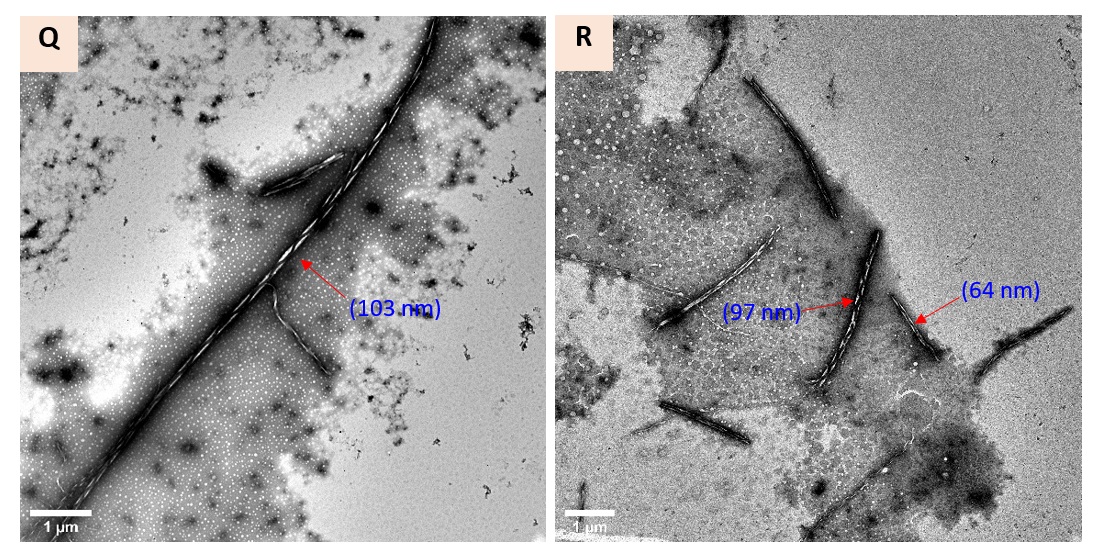

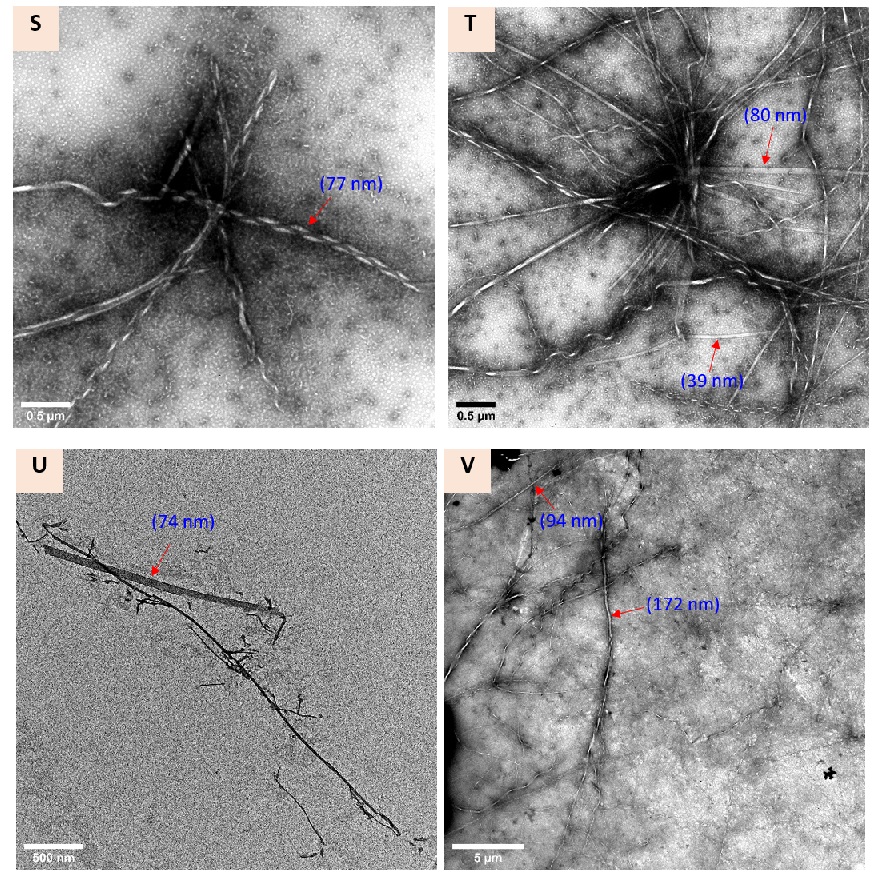

**Figure S4. Variations in structural characteristic of α-Syn fibrils in different class.** **(A-B)** Thin and curly fibrils. **(C-N)** Twisted ribbons. **(O-S)** Helical ribbons **(T-U)** Flat sheet and **(V)** Nanotube. For twisted ribbons width and period have been mentioned, respectively, for all other class width has been mentioned in the brackets.

**A**

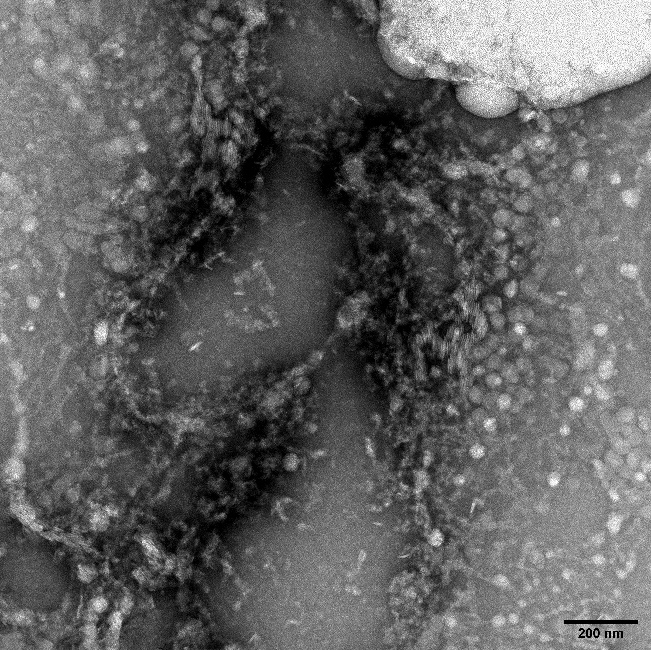

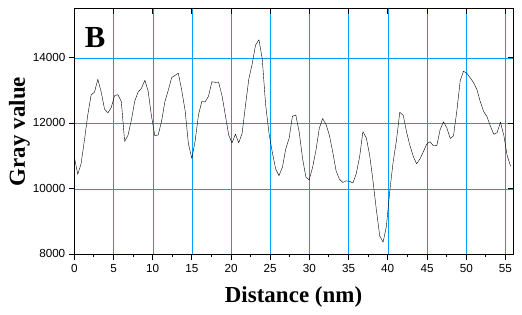

**B**

**Figure S5.** Presence of nanodiscs- like structure at early stage of α-Syn aggregation in the presence of DMPS vesicles **(A)** TEM images of nanodisc-like structure **(B)** Distance analysis of selected region from **(Fig. A)** shows these are spaced by 3-4 nm. Distance analysis was performed with image J software.

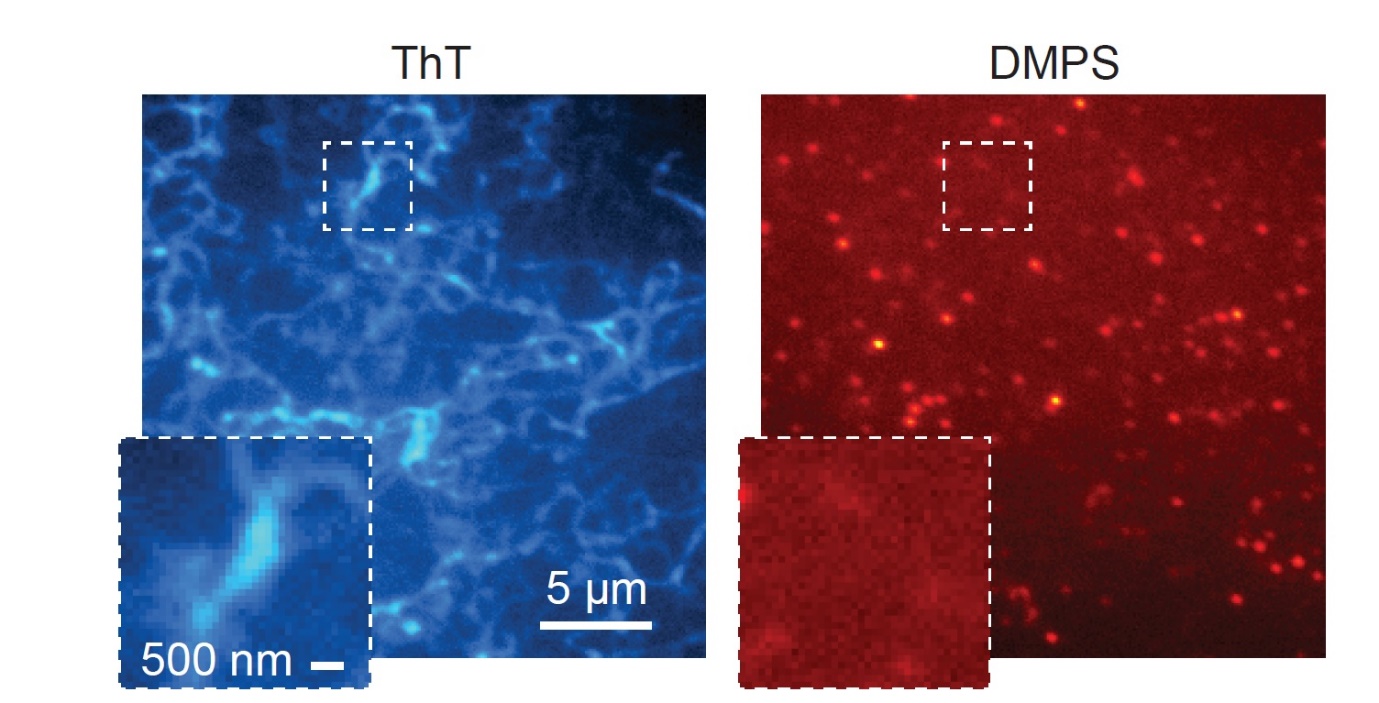

**Figure S6. TIRFM imaging. A** 0.5 µM α-syn amyloid fibrils and in the presence of 4 μM DMPS vesicles in the presence of 1 nM streptavidin-AF647 and 5μM ThT **(A)** Images were recorded for 50 frames from the red channel (AF647 emission) with 641 nm illumination, followed by **(B)** green channel (ThT emission) with 405 nm illumination.
